# Efficient and Flexible Integration of Variant Characteristics in Rare Variant Association Studies Using Integrated Nested Laplace Approximation

**DOI:** 10.1101/2020.03.12.988584

**Authors:** Hana Susak, Laura Serra-Saurina, Raquel Rabionet Janssen, Laura Domènech, Mattia Bosio, Francesc Muyas, Xavier Estivill, Georgia Escaramís, Stephan Ossowski

## Abstract

Rare variants are thought to play an important role in the etiology of complex diseases and may explain a significant fraction of the missing heritability in genetic disease studies. Next-generation sequencing facilitates the association of rare variants in coding or regulatory regions with complex diseases in large cohorts at genome-wide scale. However, rare variant association studies (RVAS) still lack power when cohorts are small to medium-sized and if genetic variation explains a small fraction of phenotypic variance. Here we present a novel Bayesian rare variant Association Test using Integrated Nested Laplace Approximation (BATI). Unlike existing RVAS tests, BATI allows integration of individual or variant-specific features as covariates, while efficiently performing inference based on full model estimation. We demonstrate that BATI outperforms established RVAS methods on realistic, semi-synthetic whole-exome sequencing cohorts, especially when using meaningful biological context, such as functional annotation. We show that BATI achieves power above 75% in scenarios in which competing tests fail to identify risk genes, e.g. when risk variants in sum explain less than 0.5% of phenotypic variance. We have integrated BATI, together with five existing RVAS tests in the ‘Rare Variant Genome Wide Association Study’ (rvGWAS) framework for data analyzed by whole-exome or whole genome sequencing. rvGWAS supports rare variant association for genes or any other biological unit such as promoters, while allowing the analysis of essential functionalities like quality control or filtering. Applying rvGWAS to a Chronic Lymphocytic Leukemia study we identified eight candidate predisposition genes, including EHMT2 and COPS7A.

**Data availability and implementation:** All relevant data are within the manuscript and pipeline implementation on https://github.com/hanasusak/rvGWAS

**Author summary:** Complex diseases are characterized by being related to genetic factors and environmental factors such as air pollution, diet etc. that together define the susceptibility of each individual to develop a given disease. Much effort has been applied to advance the knowledge of the genetic bases of such diseases, specially in the discovery of frequent genetic variants in the population increasing disease risk. However, these variants usually explain a little part of the etiology of such diseases. Previous studies have shown that rare variants, i.e. variants present in less than 1% of the population, may explain the rest of the variability related to genetic aspects of the disease.

Genome sequencing offers the opportunity to discover rare variants, but powerful statistical methods are needed to discriminate those variants that induce susceptibility to the disease. Here we have developed a powerful and flexible statistical approach for the detection of rare variants associated with a disease and we have integrated it into a computer tool that is easy and intuitive for the researchers and clinicians to use. We have shown that our approach outperformed other common statistical methods specially in a situation where these variants explain just a small part of the disease. The discovery of these rare variants will contribute to the knowledge of the molecular mechanism of complex diseases.

## Introduction

The rapidly improving yield and cost-effect ratio of Next Generation Sequencing (NGS) technologies provide the opportunity to study associations of genetic variants with complex multifactorial diseases in large cohorts at a genome-wide scale. As opposed to genome-wide association studies (GWAS), which are based on counting of genotypes at predefined genomic positions with alternative alleles of medium to high minor allele frequency in the population (MAF >1 %), whole-exome and whole-genome sequencing (WES, WGS) enable the study of rare genetic variants (RV) across the whole exome or genome, respectively. Previous studies have shown that RVs play an important role in the etiology of complex genetic diseases(1–4). Furthermore, it has been demonstrated that RVs are more likely to affect the structure, stability or function of proteins than common variants(5,6). Therefore, statistical analysis of the combined set of rare variants across genes or regulatory elements has the potential to reveal new insights into the genetic heritability of complex diseases and the predisposition to cancer. To this end, rare variant association studies (RVAS) that facilitate identification of novel disease loci based on the burden of rare and damaging variants with low to medium effect size within genomic units of interest have been developed(7).

One of the major difficulties when associating rare variants to disease is the lack of power when using traditional statistical methods like GWAS. Given that few individuals are carriers of the rare alternative allele, association studies based on single variant positions would require extremely large sample sizes. To overcome this obstacle and to increase statistical power, studies of RV consider simultaneously multiple variable positions within functional biological units, such as genes, promoters or pathways, for association to disease. Different statistical methods that address the problem of aggregated analysis of rare variants in case-control studies have been proposed. For example, score based methods pool minor alleles per unit into a measure of burden, which is used for association with a disease or phenotypic trait(8–11). These burden tests are powerful when a high proportion of RVs found in a gene affect its function and their effects on the disease are one-sided, i.e. either protective or deleterious. This is rarely the case since usually few deleterious variants coexist with many neutral and possibly some protective variants. Hence advanced methods have been developed to consider heterogeneous effects among RVs on the disease (or trait), which are mainly based on variance component tests, e.g. SKAT and C-alpha(12,13). These methods are more powerful than burden tests when the hypothesis of unidirectional effects does not hold(14). More recently, novel methods have been introduced. These contemplate that both types of genetic architectures may coexist throughout the genome, by being constructed as a linear combination between burden and variance-component tests, such as SKAT-O(15). He et al.(16) developed an alternative method, a hierarchical Bayesian multiple regression model (HBMR) additionally accounting for variant detection errors commonly produced using NGS data, by incorporation of genotype misclassification probabilities in the model. Sun et al.(17) proposed a mixed effects test (MiST) within the framework of a hierarchical model, considering biological characteristics of the variants in the statistical model. In brief, MiST assumes that individual variants are independently distributed, with the mean modeled as a function of variant characteristics and certain variance that accounts for heterogeneous variant effects. In the resulting generalized linear mixed effects model (GLMM) variant-specific effects are treated as the random part of the model and patient and variant characteristics as the fixed part. The authors claim that, under the assumption that associated variants share common characteristics such as similar impact on protein function (e.g. primarily loss of function), using this prior information increases the power of the test. However, they also note that attempting to estimate the full model for inference purposes requires multiple integration, such that it becomes too computationally intensive for a genome-wide scan. Instead, a score test under the null hypothesis of no association is proposed, avoiding multiple integration.

Building on the concept of MiST, but with the motivation of making inference based on full model estimation, we propose a Bayesian alternative to the GLMM, using the Integrated Nested Laplace Approximation (INLA) for efficient model estimation(18). Calculating the marginal likelihood to estimate complex models in a fully Bayesian manner is often infeasible. Therefore, approximate procedures such as the heuristic Markov Chain Monte Carlo (MCMC) method are conventionally applied(16). MCMC is a highly flexible approach that can be used to make inference for any Bayesian model. However, evaluating the convergence of MCMC sampling chains is not straightforward(19). Another concern with MCMC is the extensive computation time, especially in large-scale analyses such as genome-wide scans. INLA is a non-sampling based numerical approximation procedure, developed to estimate hierarchical latent Gaussian Markov random field models. Being based on numerical approaches instead of simulations renders INLA substantially faster than MCMC. Furthermore, Rue and Martino(20) demonstrated for several models that INLA is also more accurate than MCMC when given the same computational resources. The flexibility of modeling within the Bayesian framework combined with rapid inference approaches opens new possibilities for genetic association testing.

Here, we present a novel Bayesian rare variant Association Test using INLA (BATI), implemented as part of the ‘Rare Variant Genome Wide Association Study’ (rvGWAS) framework. rvGWAS combines quality control (QC), interactive filtering, detection of data stratification (technical or population based), integration of functional variant annotations and four commonly used rare variant association tests (Burden, SKAT-O, KBAC and MiST) as well as the two Bayesian alternatives, HBMR and BATI. We demonstrate using realistic benchmarks that BATI substantially outperforms existing methods if prior information on the effect of variants on protein function is used. We further show that BATI successfully copes with complex population structure and other confounders. Finally, we propose how to use ‘difference in deviance information criterion’ (ΔDIC) for model selection.

## Material and Methods

### Bayesian rare variant Association Test based on Integrated nested Laplace approximation (BATI)

Integrated Nested Laplace Approximation is a recent approach to implement Bayesian inference on latent Gaussian models, which are a versatile and flexible class of models ranging from (generalized) linear mixed models (GLMMs) to spatial and spatio-temporal models. A detailed definition of INLA can be found in(18,21,22). Here we applied INLA using the implementation of the R-INLA project (R package INLA version 17.06.20) to build a hierarchical Bayesian approach to the GLMM for the association of rare variants with phenotypes in the context of case-control studies. Our method termed BATI can efficiently and flexibly integrate a large number of categorical and numeric characteristics of genetic variants as covariates, as INLA facilitates estimation of the full model even for complex structures of random effects.

### Model specification

Assume we have N individuals, and let *Y*_*i*_ (*i* = 1,…, *N*) be the observed phenotype of the *i*th individual that belongs to an exponential family:

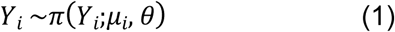

where the expected value *μ* = *E*(*Y*_*i*_) is linked to a linear predictor *η*_*i*_ through a known link function *g*(·), so that *g*(·) = *η*_*i*_. In our case *Y*_*i*_ is a binary variable representing affected individuals (cases) vs. unaffected individuals (controls). We propose to construct the likelihood of the data based on a logistic distribution and use the identity function for *g*(·). The linear predictor *η*_*i*_ is defined to account for potential confounding covariates at the individual level as well as for covariates at the variant level such as a variant’s functional impact:

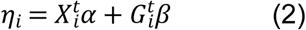

where *X*_*i*_ is a *m* × 1 vector of individual-based confounding covariates and *G*_*i*_ denotes a *p* × 1 vector of genotypes for *p* RVs. Each genotype is coded as 0, 1, or 2, representing the number of minor alleles. *α* and *β* are the regression vectors of coefficients.

BATI can account for individual variant characteristics under the assumption that similar variant-specific characteristics have a similar effect on the function of the protein and hence the phenotype, while still allowing for potential variant-specific heterogeneity effects. Thus *β* can be modeled in a hierarchical way as:

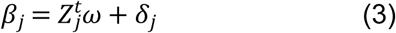

where *ω* is a vector of *q* × 1 (*j* = 1,…,*q*) variant-specific regression coefficients, *Z*^*t*^ is a *p* × *q* matrix (for *q* covariates per variant), and *δ* is a *p* × 1 random effects vector which is assumed to follow a multivariate Gaussian distribution with mean 0 and covariance matrix *τQ*. If no dependency structure is defined across variants, as in MiST(17), *Q* is a *p* × *p* identity matrix and *τ* the random effects variance. However, in order to model a correlation structure between variants, such as physical distance dependency, BATI allows to construct *Q* such that it reflects this structure. This is enabled by INLA, which provides Laplace approximation of the posterior distributions, therefore allowing the estimation of the full model for complex structures of random effects.

Plugging equation (3) into (2) we obtain the expression of a generalized linear mixed effects model (GLMM):

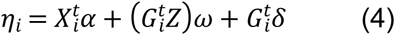

with *α* and *ω* as fixed effects coefficients and *δ* as random effects coefficients. Given the vector of parameters *θ* = {*α,ω,δ*}, the objectives of the Bayesian computation are the marginal posterior distributions for each of the elements of the parameter vector *p*(*θ*_*s*_|*y*) and for the hyper-parameter *p*(*τ*|*y*). In order to compute the marginal posterior for the parameters, we first need to compute *p*(*τ*|*y*) and *p*(*θ*_*s*_|*τ,y*). The INLA approach exploits the assumptions of the model to produce a numerical approximation to the posteriors of interest, based on the Laplace approximation(23).

### Model selection

The classical approaches of association tests are based on hypothesis testing, where the null hypothesis assumes no genetic effects, and the alternative hypothesis assumes a genetic effect on the phenotype. In the context of BATI this can be specified as follows:

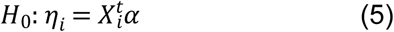

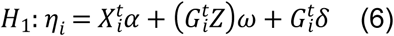

A classic Bayesian criterion for model goodness of fit is the *Deviance Information Criteria* (*DIC*)(24). *DIC* is calculated as the expectation of the deviance over the posterior distribution plus the effective number of parameters. Thus, difference in *DIC* between the H_0_ and the H_1_ models, 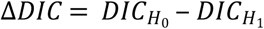, can be used as the model selection criteria. As a rule of thumb values of Δ*DIC* > 10 are recommended to reject the null-hypothesis. However, to evaluate the ability of Δ*DIC* to correctly choose between null or alternative models we suggest the use of simulations, as proposed by Holand et al.(25). To find an estimate of the probability of type I error, concluding that there are genetic effects when in truth there is none, we randomly assign individuals to either cases or controls. We then adjust models under null and alternative hypothesis for each gene or biological unit included in the genome wide study, obtaining the empirical distribution of Δ*DIC*. Finally, we select a *ΔDIC* threshold from the quantile corresponding to the desired significance level. For more robust threshold estimation, we propose to generate *S* datasets by randomly shuffling cases and controls, such that *S ΔDIC* thresholds can be obtained and the median of the thresholds can be used. We used *S* = 10 for model selection in our benchmark study.

### A comprehensive framework for rare variant association analysis (RVAS)

We developed the ‘Rare Variant Genome Wide Association Study’ (rvGWAS) framework (Fig 1A and Supplementary S1 Fig), an all-in-one tool designed for RVAS tests using case-control cohorts analyzed by NGS. rvGWAS supports rare variant association aggregating by genes or any other biological unit such as promoters or enhancers. It provides all essential steps and functionalities to perform the complete analysis of whole-exome sequencing (WES) or whole-genome sequencing (WGS) based case-control study designs: (1) it facilitates comprehensive quality control and filtering, (2) it evaluates data stratification (either technical or population based), (3) it enables the integration of patient- and/or variant-based covariates in association tests in an easy and intuitive fashion, and (4) it integrates six conceptually different rare-variant association methods. It is implemented in a modular way and provides great flexibility, allowing to analyze a wide range of association study designs.

**Fig 1.**
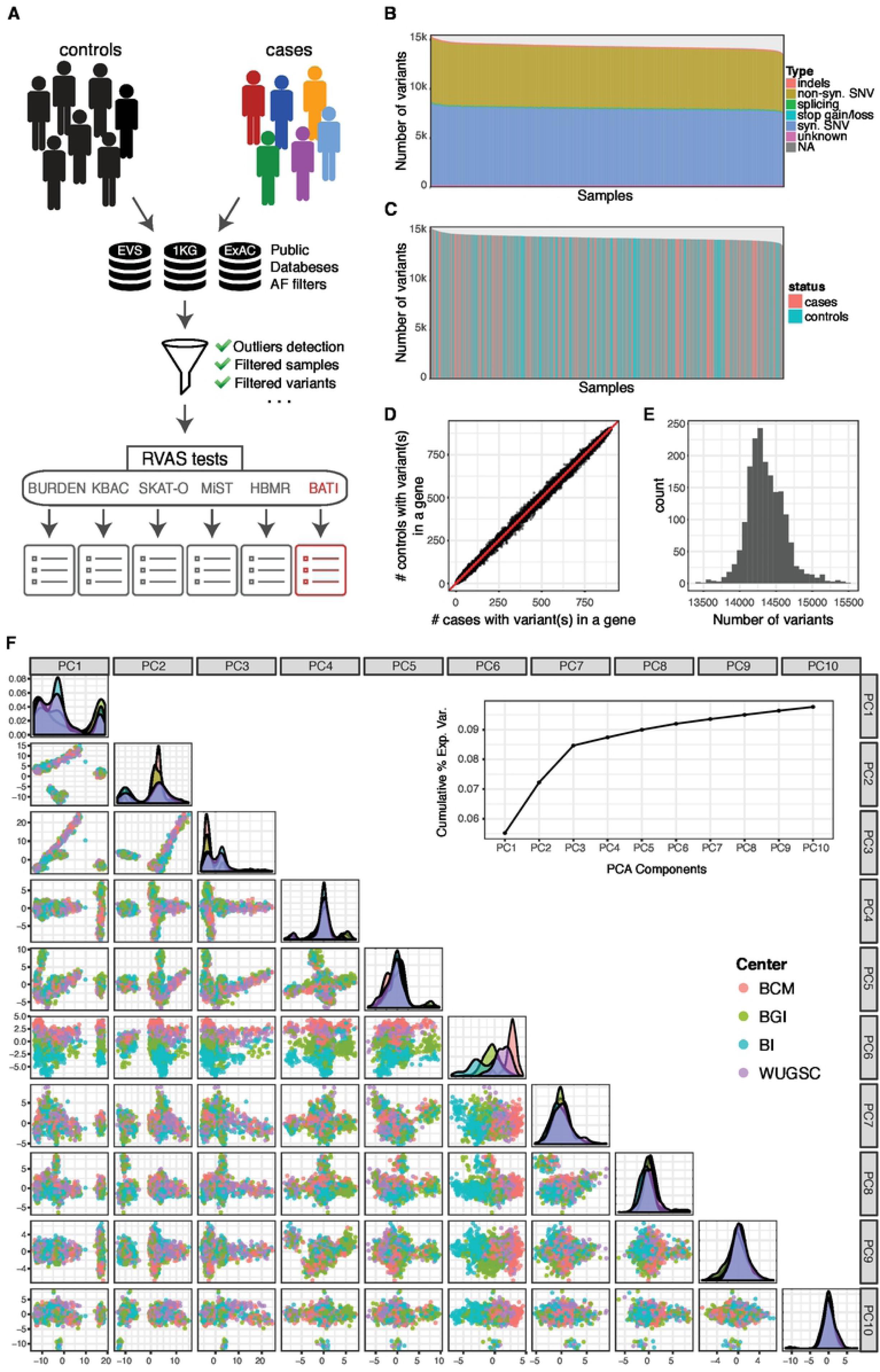
rvGWAS workflow and QC plots for 1810 high quality samples from 1000GP used for benchmarking. (**A)** rvGWAS workflow for performing QC and six RVAS tests. The QC module computes quality statistics shown in panels B-F. The result of each RVAS test is a ranked list of genes with various informative attributes. (**B)** Bar-plot for number of variants per sample, colored by functional annotation of variants. (**C)** Barplot for number of variants per sample, colored by assignment to cases (∼1/2) or controls (∼1/2). (**D)** Number of variants per gene in cases (x-axis) and controls (y-axis). Each dot is one gene, while the red line shows the ratio of the number of cases and controls (1:1). (**E)** Histogram for number of mutations per sample after removal of outliers. (**F)** Projection on first 10 PCA components. Samples are colored by sequencing center. The graph in the upper right corner shows the cumulative percentage of variance explained per principal components. Principal components can be used as covariates in several RVAS tests.

BATI and five other RVAS methods are integrated in the rvGWAS framework. KBAC, SKAT-O, and MiST, were chosen to be included due to their superior performance compared to eight other RVAS methods in a benchmark study by Moutsianas et al.(14). In addition, we included the classical Burden test representing the most simplistic and intuitive form of RVAS tests. Finally, we incorporated HBMR, which is conceptually the most similar to BATI in terms of its estimation approach (while MiST is more similar in terms of model specification). The six supported RVAS tests represent a broad spectrum of approaches, including classic aggregation of variants as a Burden variable, variance component bidirectional tests, mixed effect models and Bayesian inference.

rvGWAS is implemented as a pipeline of R scripts, and is available online at https://github.com/hanasusak/rvGWAS. Detailed descriptions of the tool, included methods as well as parameters are provided in supporting information file.

### Realistic ‘semi-synthetic’ simulations of whole-exome sequencing based case-control studies

To allow for benchmarking using highly realistic disease cohorts, which correctly represent all expected sources of noise, we developed a new disease cohort simulator combining thousands of real WES datasets from various studies with known risk variants for a selected disease type. The simulator randomly assigns WES samples to the case or control group and introduces predisposition variants found in ClinVar for a disease of choice into the VCF files of cases.

We used two large datasets as basis for the simulation: 1) WES data of the 1000 Genomes Project (1000GP), and 2) an in-house dataset combining patients diagnosed with various conditions and healthy individuals subjected to WES during 2012 to 2017. VCF files from 1000GP (phase3)(26,27) were downloaded from ftp://ftp.1000genomes.ebi.ac.uk/vol1/ftp/release/20130502/. This cohort contains 2504 individuals from 26 populations. WES libraries of 1000GP were prepared using one of four oligo enrichment kits: (1) Nimblegen SeqEz V2, (2) Nimblegen SeqEz V3, (3) VC Rome, and (4) Agilent SureSelect V2. Additional sample information used as covariates (population, super population, gender) was obtained from the file integrated_call_samples_v3.20130502.ALL.panel. We excluded related individuals, e.g in parent-child trios we included the parents (if not consanguineous), but not the child. To minimize issues with population stratification due to highly diverse populations we only included individuals not belonging to African ancestry populations, as Africans had on average 25% more variants than individuals from other ancestry groups. Nonetheless, the remaining cohort still represents a mixed population, allowing us to benchmark population stratification efficiency of the RVAS tests.

The in-house ‘Iberian’ WES cohort includes 1189 individuals of Spanish ancestry and is therefore highly homogeneous. WES libraries were prepared using three different oligo enrichment kits: (1) Agilent SureSelect 50, (2) Agilent SureSelect 71, and (3) Nimblegen SeqEz V3. Computational analysis and variant calling was performed according to GATK best practice guidelines (https://software.broadinstitute.org/gatk/best-practices/). For simulation purposes we only considered genomic loci that were targeted and covered with at least 10 sequence reads by all oligo enrichment kits, and variants with a call rate higher than 85%. Samples that were identified as outliers based on the number of called variants, transition to transversion (Ti/Tv) ratio, or their projection on the first two principal components from principal component analysis were removed from further analysis. The remaining datasets, named 1000GP and Iberian cohort, consisted of 1,810 and 1,167 samples harboring 493,314 and 285,658 unique loci with alternative alleles, respectively. From 1000GP we randomly selected half of the samples as cases, the other half as controls, while for the Iberian cohort we selected one third as cases, and two thirds as controls.

### Simulating a breast cancer risk cohort

To introduce realistic disease variants into a ‘semi-synthetic’ breast cancer predisposition cohort, we queried the ClinVar database for breast cancer risk variants annotated as exonic or splicing. We removed variants that had MAF higher than 0.01 in any ancestry population in any of three commonly used exome databases: EVS, 1000GP or ExAC. Six genes had more than five annotated disease risk variants in ClinVar: *BRCA2 (MIM: *600185), BRCA1 (MIM: *113705), PALB2 (MIM: *610355), BRIP1 (MIM: *605882), CHEK2 (MIM: +604373)* and *BARD1 (MIM: *601593)* (Supplementary S1 Table), which we used to simulate risk patients by adding variants to the VCF files (zero or one variant per case). As expected, all six genes already had rare variants, likely benign, in the unmodified cohorts (Supplementary S2 Table and Supplementary S3 Table). This type of noise is expected in any case-control study using WES data, and hence makes the simulation more realistic. We generated three genetic architectures per gene, with ∼2% (1), ∼1% (2) or ∼0.5% (3) of phenotypic variance explained (VE) by introducing ClinVar risk variants. To this end we used the method of So et al.(28) for calculation of cumulative VE each time a variant was added to a gene until the targeted VE was reached. Calculation of VE requires three parameters per each variant: the prevalence of the trait, the population frequency of the risk allele, and the genotype relative risk (RR). In practice, only odds ratios (OR) are available in many case-control studies. However, OR approximates RR when the disease prevalence in a population is low(28). As prevalence of breast cancer we selected an estimate for the Spanish population of 0.00085(29). In order to generate realistic RR distributions, we generated a distribution (Supplementary S2 Fig) assuming that the likelihood of having high RR is negatively correlated with MAF(14). For *BRCA1* and *BRCA2* we simulated two different types of genetic architectures, by introducing in one architecture only missense variants, and in the other only loss of function (LoF) SNVs (i.e. stop-gain, stop-loss or splicing). This allowed us to test if MiST and BATI benefit from features that capture biological function and context of variants. For the four remaining genes, the variants were simulated regardless of their functionality. The simulation procedure is repeated 100 times for each of the 8 architectures in order to generate 100 datasets for evaluation of statistical power and type I error rates (TIER). For *BARD1* it was not possible to reach the desired VE of 2% and 1% in most simulations due to an insufficient number of breast cancer risk variants found in ClinVar. Supplementary S3 Fig and Supplementary S4 Fig show the exact levels of VE in 100 simulations per gene for each of the two cohorts.

## Results

### Quality control and filtering of benchmark WES cohorts

Cohorts used for benchmarking of test methods consisted of 1,810 individuals in the 1000GP cohort and 1,167 individuals in the Iberian cohort, harboring 493,314 and 285,658 unique loci with a non-reference genotype in at least one of the samples, respectively. Both datasets were analyzed and filtered using the rvGWAS quality control modules (see Methods and Supporting information file). For benchmarking purposes, we only considered variants in regions targeted by all used oligo enrichment kits. However, in the case of the Iberian cohort we observed that a small subset of regions supposed to be targeted consistently showed low coverage in a kit-specific manner, leading to strong biases identified by the data stratification module of rvGWAS (data not shown). The bias disappeared when excluding regions with less than 10x average coverage in at least one kit (Supplementary S5F Fig). Samples included in the final simulation cohorts show no biases in any of the first ten components of the PCA (1000GP: Fig 1F, Iberian: Supplementary S5F Fig), and the explained variance per PCA component is low (Fig 1F, Supplementary S5C Fig). Furthermore, samples in the two cohorts show a normal distribution of the number of mutations (Fig 1E, Supplementary S5E Fig) and Ti/Tv ratio (data not shown), and show no bias in the number of variants and fractions of InDels or synonymous, nonsynonymous and LoF SNVs (Fig 1B, Fig 1C, Supplementary S5A-B Fig). Finally, there is a high correlation between the fraction of cases and of controls having variants in any given gene (Fig 1D, Supplementary S5D Fig).

### Benchmarking RVAS Tests using semi-synthetic breast cancer risk cohorts

We used the rvGWAS framework to benchmark the six RVAS tests (Burden, SKAT-O, KBAC, MiST, HBMR and BATI) on the 1000GP and Iberian cohorts with simulated breast cancer risk variants. In order to simulate a realistic breast cancer predisposition case-control study we randomly split each of the original cohorts in a case (1000GP: 905, Iberian: 389 samples) and a control group (1000GP: 905, Iberian: 778 samples), and, in the case group samples, added ClinVar risk variants to the genes *BRCA2, BRCA1, PALB2, BRIP1, CHEK2* and *BARD1* using realistic variance explained (VE) rates (see Methods). Before performing the RVAS we filtered out common variants (AF>0.01 in public databases or in the randomized control group) as well as variants that were annotated as synonymous or had a CADD score below 10 (likely benign, see https://cadd.gs.washington.edu/info). For BATI and MiST we used prior information on variant characteristics as covariates: CADD scores as a quantitative variable and exonic function (missense, loss-of-function, InDels) as a categorical variable. We repeated the simulation and benchmarking process 10 times, including the randomized case-control assignment in order to randomize background noise in each benchmark cycle.

### Type I Error Rate estimates

The six benchmarked RVAS tests use diverse criteria for statistical significance (p-value, Bayes factor or Δ*DIC*). To generate comparable significance thresholds, we performed RVAS tests on randomly split cohorts, but without introduced ClinVar risk variants. Hence, significant associations should only be found by random chance and constitute false positives. This procedure allowed us to obtain comparable thresholds for desired type I error rates for all methods. For each of the 10 random cohort splits we obtained p-value significance thresholds for Burden, KBAC, SKAT-O and MiST that translate to 5%, 0.1% and 0.01% TIER. Similarly, for HBMR and for BATI we calculated thresholds for Bayes factor and Δ*DIC* resulting in the same TIER levels. Estimated thresholds are highly similar across all 10 randomized case-control splits (Supplementary S6 Fig). At 0.01% TIER only 2 genes (out of ∼20,000) are expected as significant by chance, therefore the observed small fluctuation of estimated significance thresholds is not surprising. We finally used the test-specific median from 10 random splits as thresholds to label a gene as significant for subsequent power analyses (Supplementary S6 Fig, Table 1 and Supplementary S4 Table).

**Table 1.** P-value, Bayes Factor (HBMR) and ΔDIC (BATI) thresholds for Type I error rates (TIER) of 0.05, 0.001 and 1e-04 estimated on 1000GP. We randomly permuted case and control labels 10 times and for each estimated empirical thresholds for each RVAS test. The median TIER values from 10 random permutations are used as thresholds for benchmark comparison.

| Method | 0.05 TIER | 0.001 TIER | 1e-04 TIER |
| --- | --- | --- | --- |
| BURDEN | 0.0519 | 1.12e-03 | 7.79e-05 |
| KBAC | 0.0650 | 1.52e-03 | 1.52e-04 |
| SKAT-O | 0.0563 | 1.47e-03 | 1.66e-04 |
| MiST | 0.0766 | 2.26e-09 | 3.33e-16 |
| HBMR | 1.2678 | 3.5774 | 9.0838 |
| BATI | 2.3898 | 9.5929 | 14.4623 |

We noticed that MiST shows zero inflated p-values (Supplementary S7A Fig). These unexpected zero p-values occur exclusively for genes with few variants (<10) across the cohort, indicating that the MiST method fails to obtain accurate p-values for genes with low burden of variants. Hence, we removed all genes with p-value 0 from MiST results (Supplementary S7B Fig). No other method showed a p-value inflation artefact or unexpectedly high Bayes Factor or ΔDIC values (Supplementary S7C-G Fig).

### Power analysis for six RVAS test methods

We next determined the power of the competing RVAS tests to identify the 8 breast cancer risk genes (*BRCA1-Missense, BRCA1-LoF, BRCA2-Missense, BRCA2-LoF, PALB2, BRIP1, CHEK2* and *BARD1*) at the three TIER levels 5%, 0.1% and 0.01% and at three levels of VE of 2%, 1% and 0.5% (1000GP: Fig 2, Iberian: Supplementary S8 Fig). For the 1000GP cohort we found that all methods showed a power close to 100% at a TIER of 5% across all tested VE levels, except for Burden and KBAC, which showed decreased performance for VE = 0.5% (Fig 2A-C left). Testing 20,000 genes (whole exome) at a TIER of 5% we expect around 1000 false positive genes, which is a poor choice for most studies. Using a TIER of 0.1% (∼20 false positive genes expected), differences between the tests become more pronounced, with Burden, KBAC and MiST showing decreased power already for 1% VE, and all methods except for BATI showing decreased power at 0.5% VE (Fig 2A-C middle). Interestingly, Burden, KBAC and SKAT-O show strongly fluctuating power for the 8 tested genes, often showing either 100% or 0% power (Fig 2C middle), meaning a risk gene was either identified in all 100 simulations, or in none. BATI achieved more than 75% power for all genes, with a median above 90%. Using a strict TIER of 0.01% (2 false positives expected for the whole exome), all tools except for MiST are able to identify risk genes at 2% VE at almost 100% (for the outlier *BARD1* we did not achieve 2% VE in all simulations due to a lack of variants in ClinVar). However, performance of all methods except BATI drops substantially for 1% VE. At 0.5% VE most methods miss the majority of risk genes in the majority of simulations (median power close to zero), while BATI still achieves a median power of 60% (Fig 2A-C right). Note that MiST performed very poorly for the strict TIER thresholds of 0.1% and 0.01%, likely due to the aforementioned zero-p-value inflation issue, which results in a large number of false positives.

**Fig 2.**
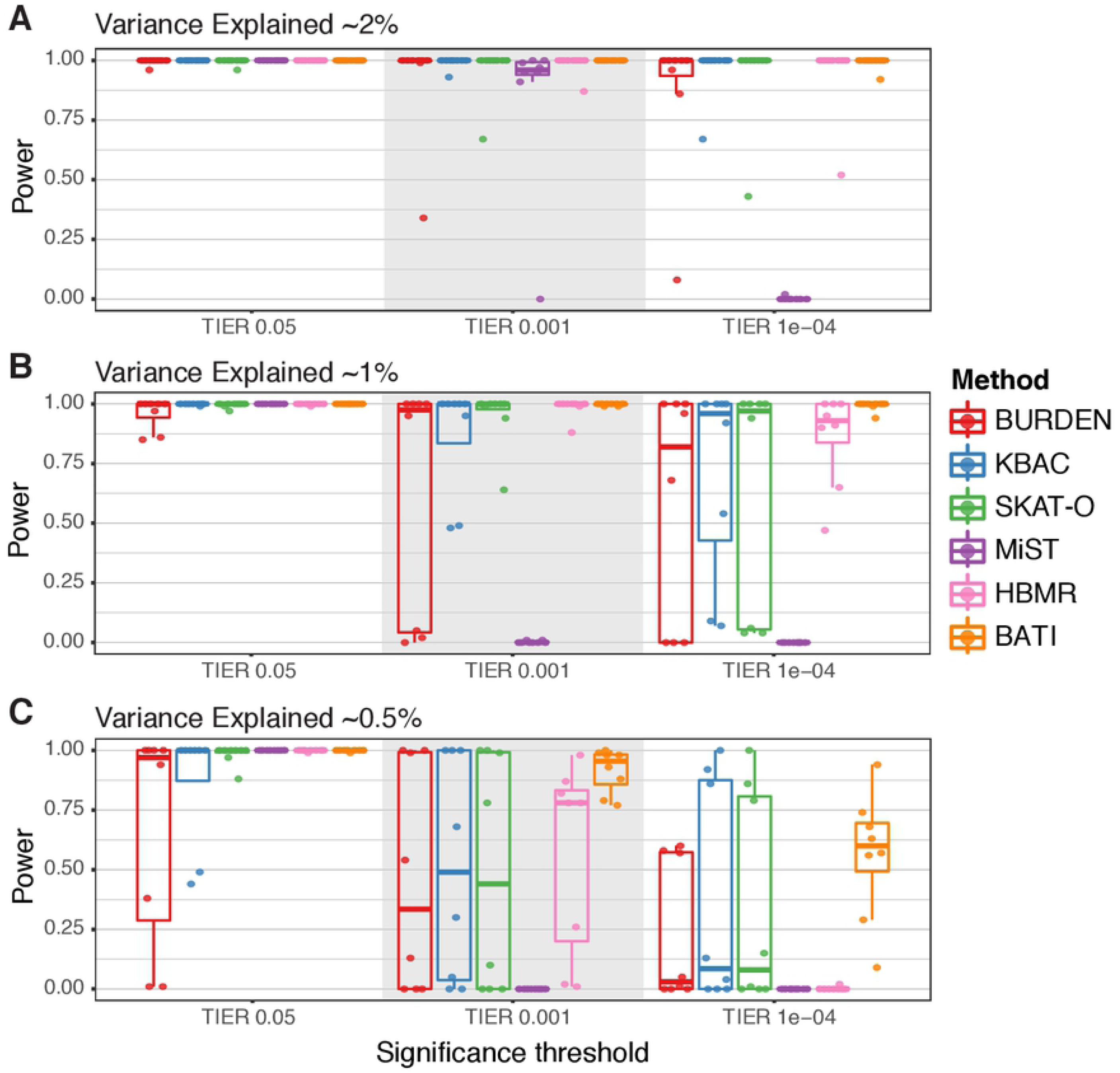
Benchmarking power of RVAS methods for the 1000GP-based BRCA risk study. Each dot in the plots represents one of simulated 8 risk genes, and y-axis values show the fraction of 100 simulations in which the gene was called as significant. RVAS tests were benchmarked under the following 9 settings. Variance explained (VE) of the incorporated risk variants is (A) ∼2%, (B) ∼1%, and (C) ∼0.5%. For each VE we tested three TIER levels, left: TIER 5%, middle: TIER 0.1%, and right: TIER 0.01%.

Results are mostly similar in the benchmark using the Iberian cohort (Supplementary S8 Fig). However, most tests perform slightly worse due to the smaller size of the cohort (1,167 vs 1,810 total individuals). Notably, BATI’s performance is stable despite the smaller cohort size. Specifically, for a low VE of 0.5% and a strict TIER of 0.01% (Supplementary S8 Fig right), all methods except for BATI show power close to 0, while BATI achieves power close to 100% for three risk genes (median power of 55%).

### Risk gene-wise power analysis

Each gene has a different architecture, i.e. rate of (likely benign) rare variants in the original cohorts, functional impact estimates for known risk variants, fraction of stop-gain or splicing variants etc. We therefore benchmarked the performance of all RVAS tests across 100 simulations of risk variants for each gene separately (1000GP cohort: Fig 3 and Table 2, Iberian cohort: Supplementary S9 Fig). In the gene-wise power plots we indicate the three TIER thresholds using red (5%), green (0.1%) and blue (0.01%) lines. Note that due to different y-Axis scaling these lines are not on the same height for different tests. As expected all methods except MiST identify all risk genes at 0.01% TIER in the 2% VE setting. However, substantial differences in power of the tests appear when VE is only 1% or 0.5%. While BATI calls most genes with TIER 0.01% even at VE of 0.5%, and all genes at TIER 0.1% with >80% power (Table 2), Burden, KBAC and SKAT-O recurrently fail to call *BRCA2* (both missense and LoF versions), and HBMR fails to call *BARD1, CHEK2* and *PALB2* already at TIER 0.1% (Table 2). The performance of Burden, KBAC and SKAT-O varies considerably between genes, while MiST, HBMR and BATI show relatively small differences. Interestingly, the power plots at 0.5% VE look very similar when comparing Burden, KBAC and SKAT-O, indicating that these methods share the same strengths and weaknesses.

**Table 2.** Power of six RVAS methods for 8 genes/architectures simulated using the 1000GP cohort and ClinVar disease variants. 100 Architectures were simulated for each gene. For BRCA1 and BRCA1 simulation was performed in missense and in LoF mode (see Methods). Power is shown for VE = 0.05% and TIER levels 0. 001 and 1e-04.

| Gene<br>Method | BRCA1<br>MiSS | BRCA1<br>LoF | BRCA2<br>MiSS | BRCA2<br>LoF | BARD1 | BRIP1 | CHEK2 | PALB2 |  |
| --- | --- | --- | --- | --- | --- | --- | --- | --- | --- |
| BURDEN | 99 | **100** | 0 | 0 | 0 | **100** | 13 | 54 | TIER=<br>0.001 |
| KBAC | **100** | **100** | 0 | 0 | 5 | **100** | 30 | 68 |  |
| SKAT-O | 99 | **100** | 0 | 0 | 0 | **100** | 10 | 78 |  |
| MiST | 0 | 0 | 0 | 0 | 0 | 0 | 0 | 0 |  |
| HBMR | 78 | 78 | 87 | 82 | 2 | 98 | 26 | 1 |  |
| BATI | 98 | **100** | **88** | **99** | **79** | 98 | **77** | **93** |  |
| BURDEN | 57 | 60 | 0 | 0 | 0 | 58 | 1 | 5 | TIER=<br>1e-04 |
| KBAC | **86** | 92 | 0 | 0 | 0 | **100** | 4 | 13 |  |
| SKAT-O | 79 | 86 | 0 | 0 | 0 | **100** | 1 | 15 |  |
| MiST | 0 | 0 | 0 | 0 | 0 | 0 | 0 | 0 |  |
| HBMR | 0 | 0 | 0 | 0 | 0 | 0 | 2 | 0 |  |
| BATI | 63 | **94** | **56** | **74** | **9** | 68 | **29** | **57** |  |

**Fig 3.**
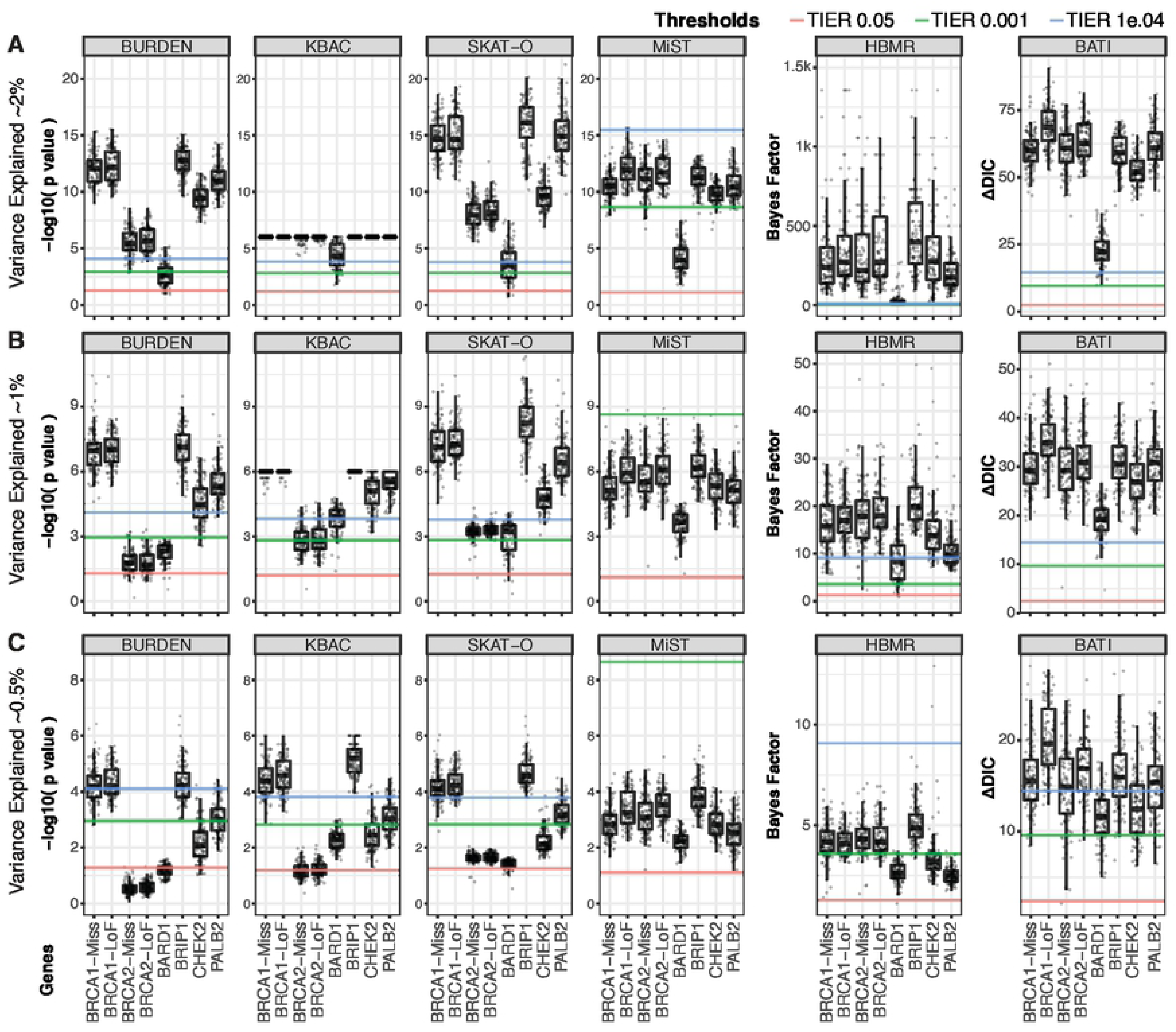
Benchmarking statistical power to detect rare variant associations for 8 genes individually. Rare variants annotated for increased breast cancer risk were simulated into the 1000GP dataset with cases and controls randomly assigned. Power (y-axis) per gene for 6 methods (Burden, KBAC, SKAT-O, MiST, HBMR and BATI) is shown for (A) 2%, (B) 1%, and (C) 0.5% variance explained between cases and healthy controls. (Due to using real SNVs in the simulation the variance explained per gene fluctuates slightly around the targeted VE. See Supplementary S3 Fig). Lower, middle and upper lines indicate relaxed (5%), medium (0.1%) and strict (0.01%) TIER thresholds, respectively.

Only MiST and BATI are able to leverage categorical variant characteristics, here represented as functional annotations such as ‘missense’, ‘LoF’, ‘indel’. As background LoF variants are rare we expected that both methods excel at predicting *BRCA1* and *BRCA2* under the LoF-architecture simulation. Indeed, for both methods we see a better performance for *BRCA1*-LoF and *BRCA2*-LoF compared to the *BRCA1*-missense and *BRCA2*-missense, respectively. For BATI, this difference is significant for both genes (*BRCA1*: p = 4.0e-13 and *BRCA2*: p = 0.0025 for VE = 0.5 using Wilcoxon rank test). As a result, BATI predicts *BRCA2*-LoF at the highest significance level (TIER 0.01%), while all other methods perform poorly. *BRCA1*-LoF shows the highest ΔDIC value from all 8 risk genes, demonstrating that the BATI method strongly benefits from categorical functional annotations.

The strong performance of BATI in terms of precision and recall comes at the price of longer run time (Supplementary S5 Table). Inference based on full model estimation leads to a higher computational complexity and hence higher run time of BATI compared to all competing methods. The computational time and complexity of RVAS test methods is a concern, as exome and genome sequencing datasets have been increasing dramatically in sample size recently. However, the INLA implementation used by BATI (R-INLA project) facilitates the use of multiple cores, and scales close to linearly with the number of used cores, allowing for analysis of large cohorts on modern servers with many cores. Moreover, lowering the allele frequency threshold of included rare variants (e.g. from AF <= 1% to AF <= 0.1%) for very large cohorts can dramatically reduce computation times.

### RVAS of chronic lymphocytic leukemia identifies candidate risk genes

Chronic lymphocytic leukemia (CLL) is a cancer of B-lymphocytes, which expands in the bone marrow, lymph nodes, spleen and blood. With the aim to identify the landscape of germline risk genes that can predispose an individual to CLL, we applied BATI and the other five competing RVAS methods integrated in rvGWAS. The CLL cohort of 436 cases was collected and sequenced following the guidelines of the International Cancer Genome Consortium (ICGC)(30) within the framework of the Spanish ICGC-CLL consortium(31) (Puente *et al.* 2015). In addition, 725 individuals from our Iberian cohort were used as controls. For the gene-wise RVAS test we preselected rare (MAF≤ 0.01 in our control cohort, ExAC and 1000GP) and potentially damaging variants (CADD score > 10). All RVAS methods were adjusted for the first 10 principal components to account for population stratification and technical biases. For BATI and MiST we additionally added the exonic function of the variants (i.e. LoF, missense, indel) and the CADD damage score as covariates. We tested all genes with a variant call rate of at least 95% and removed genes flagged by Allele Balance Bias (ABB)(32) as enriched with false positive variant calls (see Supporting information file for details). BATI identified 12 candidates that passed the significance threshold of 10^−4^ (Supplementary S6 Table). Among those, EHMT2 and COPS7A are promising CLL risk gene candidates. The heterodimeric methyltransferases EHMT1 and EHMT2 have recently been implicated with prognosis of CLL and CLL cell viability(33). COPS7A (previous name COP9) is involved in the Transcription-Coupled Nucleotide Excision Repair (TC-NER) pathway and the COP9 signalosome complex (CSN) is involved in phosphorylation of p53/TP53, JUN, I-kappa-B-alpha/NFKBIA, ITPK1 and IRF8/ICSBP. However, replication of results in independent cohorts is required to evaluate these findings.

## Discussion

Here we presented a comprehensive framework, rvGWAS, to facilitate user-friendly and intuitive analysis of RVAS in case-control studies using whole genome or custom-captured next generation sequencing data. rvGWAS integrates data quality control and filtering, several existing rare variant association tests and the newly developed BATI test. We showed how BATI leverages both categorical and numerical variant characteristics and strongly benefits from their inclusion as covariates. We demonstrated BATI’s significant gain in power if risk genes contain mostly LoF variants, while still performing at least as good as other methods when testing genes containing mostly missense variants.

Model estimation when using complex data structures, including exome-wide genetic variants, numerical damage estimates and functional annotations, becomes computationally heavy. Therefore, existing tests do not estimate the full model (as in MiST) or use the relatively slow MCMC (as in HBMR). BATI addresses this issue by estimating the full model using Integrated Nested Laplace Approximation, which requires reasonable computational resources even when using complex data structures. INLA provides approximations to the posterior marginals of the latent variables, which are accurate and extremely fast to compute(18). INLA was originally developed as a computationally efficient alternative to MCMC and presents two major advantages. On the one hand, INLA’s fast speed allows it to work on models with huge dimensional latent fields and a large number of covariates at different hierarchical levels (for example in case of RVAS at the patient level and at the variant level). On the other hand, INLA treats latent Gaussian models in a unified way, thus allowing for greater automation of the inference process. Thanks to these characteristics, INLA has already been used in a great variety of applications(34–39). Leveraging the efficiency of INLA, BATI, unlike MiST, can make inference based on full model estimation, and provides comprehensive information on estimates of model parameters. Furthermore, BATI allows for the inclusion of many numerical or categorical features as covariates. Which other features, in addition to functional impact and functional annotation of variants, could be beneficial for association testing remains to be determined. Promising categories include variant call quality, tissue-specific gene expression measures, biological pathways or copy number variants.

Previous benchmark studies of RVAS tests typically relied on pure simulations of variants, for instance based on HapMap statistics, resulting in completely artificial cohorts(14). Furthermore, simulations were often restricted to small regions of the genome, limiting their power for benchmarking exome-wide association tests. Simulated variant data is well-known to lack the complexity and noise-level of real data, resulting in overly optimistic benchmark performances and unrealistic expectations of the clinical researchers. Moreover, the use of random ‘causal’ variants hampers the benchmarking of methods that leverage characteristics of causal disease variants, which are enriched in high damage scores and high impact changes such as LoF variants. Here we combined real WES cohorts, representing realistic background noise, with real disease variants, featuring realistic functional impact profiles and variant distributions, to form semi-synthetic benchmark cohorts. We developed sampling methods allowing to test different disease architectures featuring various levels of variance explained in multiple risk genes. Furthermore, tests in the original randomized cohorts without introduced disease variants facilitated the translation of method-specific significance thresholds to comparable thresholds for type I error rates.

Using these simulations, we show that methods vary substantially in power, especially for risk genes explaining a small fraction of the variance in a cohort. We found that differences between methods when VE is low (1% and 0.5%) are substantially more profound than previously appreciated, with some methods showing strongly fluctuating success rates for different genes and close to zero power at VE of 0.5%. For example, MiST showed favorable results on purely artificial benchmark sets(14), but performed poorly on our realistic WES cohorts, likely due to an issue with zero-inflated p-values caused by inappropriate handling of low variant counts. Specifically, MiST failed to identify any risk gene at low VE or low TIER thresholds. We further found that the performance patterns of Burden, KBAC and SKAT-O across the 8 risk gene architectures are highly similar when compared to MiST, HBMR and BATI. Burden, KBAC and SKAT-O fail to predict the same genes at 0.5% VE, namely BRCA2, BARD1 and CHEK2, which are characterized by high numbers of benign background variants. It is therefore likely beneficial to combine Burden- and SKAT-type methods with completely different approaches to compensate for Burden and SKAT specific weaknesses.

In summary, leveraging variant characteristics and using the fast and accurate INLA model estimation, BATI outperforms existing RVAS test methods on realistic WES cohorts using real disease variants in 8 breast cancer risk genes, in hundreds of permutations. By facilitating integration of large numbers of covariates, BATI represents a flexible testing approach that can be further extended and enhanced in the future.

## Supporting information

**S1 Text**. Supporting information for Efficient and Flexible Integration of Variant Characteristics in Rare Variant Association Studies Using Integrated Nested Laplace Approximation

